# The Timing of Cortical Activation in Associator Grapheme-Colour Synaesthetes using MEG

**DOI:** 10.1101/2022.12.16.520766

**Authors:** Giorgos Michalareas, Flor Kusnir, Gregor Thut, Joachim Gross

## Abstract

Grapheme-colour synaesthetes experience an anomalous form of perception in which graphemes systematically induce specific colour concurrents in their mind’s eye (“associator” type). Although grapheme-colour synaesthesia has been well characterized behaviourally, its neural mechanisms remain largely unresolved. There are currently several competing models, which can primarily be distinguished according to the anatomical and temporal predictions of synaesthesia-inducing neural activity. The first main model (Cross-Activation/ Cascaded Cross-Tuning) posits *early* recruitment of occipital colour areas in the initial feed-forward sweep of brain activity. The second (Disinhibited Feedback) posits: (i) later involvement of a multisensory convergence zone (for example, in parietal cortices) after graphemes have been processed in their entirety; and (ii) subsequent feedback to early visual areas (i.e., occipital colour areas). In this study, we examine both the timing and anatomical correlates of associator grapheme-colour synaesthetes (n=6) using MEG. Using unbiased analysis methods with little *a priori* assumptions, we applied Independent Component Analysis (ICA) on a single-subject level to identify the dominant patterns of the induced, synaesthetic percept. We observed evoked activity that significantly dissociates between synaesthesia-inducing and non-inducing conditions at approximately 190 ms following grapheme presentation. This effect is present in grapheme-colour synaesthetes, but not in matched controls, and exhibits an occipito-parietal topology localised consistently across individuals to extrastriate visual cortices and to superior parietal lobes. Due to the observed timing of this evoked activity and its localisation, our results support a model predicting relatively late synaesthesia-inducing activity, more akin to the Disinhibited Feedback model. In light of previous findings, our study suggest that differential timing and neural mechanisms may account for associator grapheme-colour synaesthetes, as compared to projectors.

**Highlights:**

- first MEG study to-date examining associator grapheme-colour synaesthetes
- we use single-subj. ICA to dissociate synaesthesia-inducing vs. non-inducing activity
- we observe evoked activity ∼190 ms following grapheme presentation
effects localize consistently to extrastriate visual and superior parietal cortices
- the observed timing is consistent with a model of synaesthesia supporting relatively late activity of synaesthesia-inducing stimuli
- our findings suggest that differential mechanisms may underlie different sub-types of grapheme-colour synaesthesia

## 1. Introduction

In synaesthesia, stimulation of one sensory modality triggers a perceptual or cognitive experience in another sensory modality. One of the most prominent and best-studied forms of synaesthesia is grapheme-colour synaesthesia, in which graphemes elicit specific colour percepts either in external space (termed “projector” synaesthesia) or in the mind’s eye (“associator” synaesthesia). These additional colour experiences are elicited automatically, involuntarily, and systematically in response to specific graphemes (i.e., in unique grapheme-colour pairs). Despite the perceptual reality of these induced percepts, they are not generally confused with components of the external world. This suggests that induced synaesthetic colours are not equivalent to real colour perception and may thus involve a distinct network of brain areas other than those implicated in real colour perception.

Although there have been quite a number of neuroimaging studies addressing the neural correlates of grapheme-colour synaesthesia, these have yielded conflicting and often ambiguous results (see (Hupe, Bordier, & Dojat, 2012) and (Hupe & Dojat, 2015) for critical reviews regarding the inconsistencies reported across previous studies). The bulk of these studies have taken a fMRI-based approach (Elias, Saucier, Hardie, & Sarty, 2003; Hubbard, Arman, Ramachandran, & Boynton, 2005; Hupe et al., 2012; Rich et al., 2006; Rouw & Scholte, 2010; Sperling, Prvulovic, Linden, Singer, & Stirn, 2006; Weiss, Zilles, & Fink, 2005; Zeki & Marini, 1998)for a review, see also Rouw, Scholte, and Colizoli (2011)) and have centred their debate on whether synaesthetically triggered sensations generate activation of colour areas (typically hV4). Fewer studies have employed electrophysiology, only three of which used MEG (Teichmann et al., 2021; Brang, Hubbard, Coulson, Huang, & Ramachandran, 2010; van Leeuwen, Hagoort, & Handel, 2013) rather than EEG (Beeli, Esslen, & Jancke, 2005; Brang, Edwards, Ramachandran, & Coulson, 2008; Brang, Kanai, Ramachandran, & Coulson, 2011; Jancke, Beeli, Eulig, & Hanggi, 2009; Sagiv & Ward, 2006; van Leeuwen et al., 2013; van Leeuwen, Petersson, & Hagoort, 2010; Volberg, Karmann, Birkner, & Greenlee, 2013).

The current state of affairs thus leaves the neural correlates of grapheme-colour synaesthesia unresolved. There are several models describing the proposed underlying neural mechanisms of synaesthesia (Lalwani & Brang, 2019; Hubbard, Brang, & Ramachandran, 2011). These models (e.g., Disinhibited Feedback, Cross-Activation, Two-Stage, Stochastic Resonance) primarily diverge on the predicted brain areas and timing of the induced synaesthetic activity. In this study, we use magnetoencephalography (MEG) to directly examine induced synaesthetic activity in order to advance our knowledge of the proposed mechanisms of synaesthesia. MEG serves as a happy medium between other neuroimaging techniques (i.e., fMRI and EEG), as it exhibits precise temporal resolution together with adequate sensitivity to the anatomical correlates of the measured signal.

Only two MEG studies to-date (Teichmann et al., 2021; Brang et al., 2010) have examined the induced synaesthetic percept via the presentation of achromatic, synaesthesia-inducing graphemes. Brang and colleagues (2010) showed activity in pre-defined area hV4 peaking between 111-130 ms, only 5-ms after the onset of activity in grapheme areas. These findings support a quick, direct communication between the involved brain areas (i.e., Cross-Activation or CCT models). However, the four grapheme-colour synaesthetes who were included in this study were strong *projector* sub-types. In contrast, Teichmann and colleagues (2021) used classification models on a larger group of grapheme-colour synaesthetes to examine the timing of induced colours; they distinguished between two possible induced colours evoked by achromatic graphemes and reported a later time period (around 200 ms) as corresponding to the induced synaesthetic activity. It remains unclear which anatomical areas underlie their effect, and, moreover, whether these results may generalise to the *associator* sub-types, which comprise the majority of grapheme-colour synaesthetes (Dixon, Smilek & Merikle, 2004).

We here conducted an MEG study on the time-course of neuronal responses to achromatic synaesthesia-inducing (versus non-inducing) graphemes (in analogy to Brang et al., 2010), but in six *associator* grapheme-colour synaesthetes. Due to strong inter-individual differences across associator synaesthetes, and to previous large-scale critiques on the use of strong *a priori* analytic parameters in synaesthesia studies, we opted to employ an unbiased, single-subject approach without prior assumptions (i.e., no pre-defined ROIs). To decompose the MEG signal and identify the dominant patterns of activity on a single-subject level, we used Independent Component Analysis (ICA) and cluster-based statistics (both on single-subject levels). ICA is a data-driven, blind source separation method that extracts statistically independent sources that have been mixed in the surface signal (Makeig, Debener, Onton, & Delorme, 2004; Makeig, Jung, Bell, Ghahremani, & Sejnowski, 1997). It makes no prior assumptions about the spatial locations of the combined signal. ICA is routinely used in the analysis of MEG data (Brookes et al., 2012; Brookes et al., 2011; Capilla, Belin, & Gross, 2013; Spadone, de Pasquale, Mantini, & Della Penna, 2012; Vigario, Sarela, Jousmaki, Hamalainen, & Oja, 2000). Since the neural mechanisms and the cortical areas underlying the triggered, synaesthetic percept are still very much under debate, and since additionally no “synaesthetic” event-related components have been defined, single-subject ICA is an ideal method for extracting the patterns consistently present across trials in grapheme-colour synaesthetes, including “weaker” patterns that may not necessarily manifest in the event-related averages of the raw, sensor time-series.

Our results show an absence of early, visually evoked (extrastriate) activity in associator grapheme-colour synaesthetes (in response to synaesthesia-inducing letters vs. non-inducing pseudoletters). Instead, we find evoked activity dissociating inducing vs. non-inducing graphemes (in synaesthetes but not matched controls) to peak at approximately 190 ms and exhibiting an occipito-parietal topology localised consistently across individual synaesthetes to extrastriate visual cortex and the superior parietal lobes. This is, to the best of our knowledge, the first MEG study to date investigating grapheme-colour associator synaesthesia, providing evidence for a relatively late timing of the induced synaesthetic activity, more akin to a Disinhibited Feedback model or its variants.

## 2. Materials and Methods

All experiments were conducted in accordance with the ethical guidelines established by the Declaration of Helsinki, 1994, and were approved by the local ethical committee of the College of Science and Engineering, University of Glasgow. All participants gave their written informed consent prior to inclusion in the study. All participants had normal or corrected-to-normal vision, including normal colour vision.

### 2.1 Participants

Six grapheme-colour synaesthetes (age range: 19-34, all female, all right-handed), and six controls (age range: 21-35, m/f=1/5, all right -handed) matched on age, handedness and educational level participated in this experiment. Developmental synaesthesia was established by means of two questionnaires: in the first, participants rated *statements* describing aspects of their synaesthetic experiences and provided accompanying written explanations of these (questionnaire adapted from Banissy et al. (2009)), while in the second, they rated visual *illustrations* portraying their synaesthetic experiences (questionnaire adapted from Skelton et al. (2009)) and also provided short accompanying statements describing additional aspects of their synaesthetic experiences (see Supplementary Material for copies of both questionnaires). Based on both questionnaires, all six grapheme-colour synaesthetes were classified as associators according to the projector-associator distinction (Dixon, Smilek, & Merikle, 2004), but see (Eagleman, 2012). In addition, grapheme-colour synaesthesia was tested and confirmed in all six individuals by means of a Consistency Test, which includes a surprise re-test (see Section, *Consistency Test*). At the conclusion of the study, all six controls were also screened for synaesthesia using the same written questionnaires administered to synaesthetes.

### 2.2 Consistency Test

The aim of the Consistency Test was to confirm grapheme-colour synaesthesia in all six synaesthetic participants using a test-retest reliability protocol, and to define synaesthetically inducing and non-inducing stimuli for the subsequent MEG task.

To this end, we used a computerized protocol adapted from Eagleman, Kagan, Nelson, Sagaram, and Sarma (2007), also providing normative data. Each trial began with the presentation of an achromatic grapheme (black on a medium grey background), together with a colour palette consisting of more than sixty-five thousand colours. Participants were instructed to select the colour that most closely matched their synaesthetic percept of the presented grapheme (or a “no colour” option if they lacked a colour experience for that grapheme). Participants were instructed to take their time and to be as precise as possible. Upon selection of a colour, the corresponding RGB value was automatically recorded and the next trial began. In total, there were 150 trials, corresponding to the full set of graphemes A-Z (26 total), the digits from 0-9 (10 total), and fourteen pseudo-letters (14 total) (see Figure 1), each repeated three times in randomised order. Matlab 2007b (The MathWorks, Inc.) was used to control both stimulus presentation and data collection.

**Figure 1.**
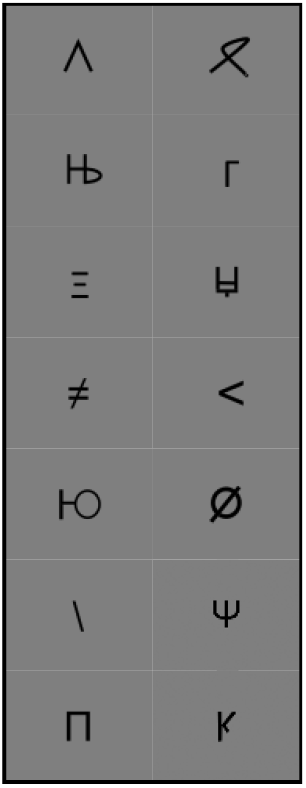
Pseudo-letters presented to grapheme-colour synaesthetes in Consistency Task. These were manually created using component features similar to letters.

After a minimum delay of three weeks, all six participants were re-tested in the exact same task. All grapheme-colour pairings were then tested for consistency, per synaesthete and across the two sessions based on the formula established by Eagleman et al. (2007): for each of the fifty graphemes (26 letters, 10 numbers, 14 pseudoletters), the total distance between the selected colours (i.e., three colours per testing session) was calculated in normalised RGB colour space. Then, all colour distances were averaged within sessions (i.e., average of fifty colour distances for the first session, and average of fifty colour distances for the second) to yield Consistency Scores for each session; and subsequently the colour distances in both sessions together (i.e., data was concatenated across sessions) were averaged to yield a Consistency Score across sessions. All six grapheme-colour synaesthetes fell within the normative synaesthesia range provided by Eagleman et al. (2007), i.e., exhibiting Consistency Scores below 1 (range of scores across sessions: 0.55-0.88).

### 2.3 Grapheme Stimuli

The set of stimuli was individually tailored to each synaesthete. First, seven colour-inducing letters and seven non-inducing pseudoletters were chosen for each synaesthete based on individual responses in the Consistency Test (i.e., those letters showing maximum consistency scores within and across sessions, and those pseudoletters showing no induced colours within and across sessions). Then, each of the seven colour-inducing letters was paired to one of the seven non-inducing pseudoletters, resulting in seven pairs of inducing/non-inducing graphemes (see Figure 2, Morph Levels 1 and 5). A sequence of “morphed graphemes” was created for each of these seven pairs, such that each of the seven stimulus sets consisted of one colour-inducing letter, one non-inducing pseudoletter, and three intermediate morphed graphemes, each representing a step-wise transformation between the preceding and succeeding grapheme pairs. These intermediate morphed graphemes were created such that they physically resembled a “blend” of the adjacent graphemes. This led to a total of thirty-five graphemes (7 stimulus sets x 5 graphemes per stimulus set) per synaesthete (also shown to individually age-matched controls). All graphemes were created manually, and were achromatic set against a medium grey background.

**Figure 2.**
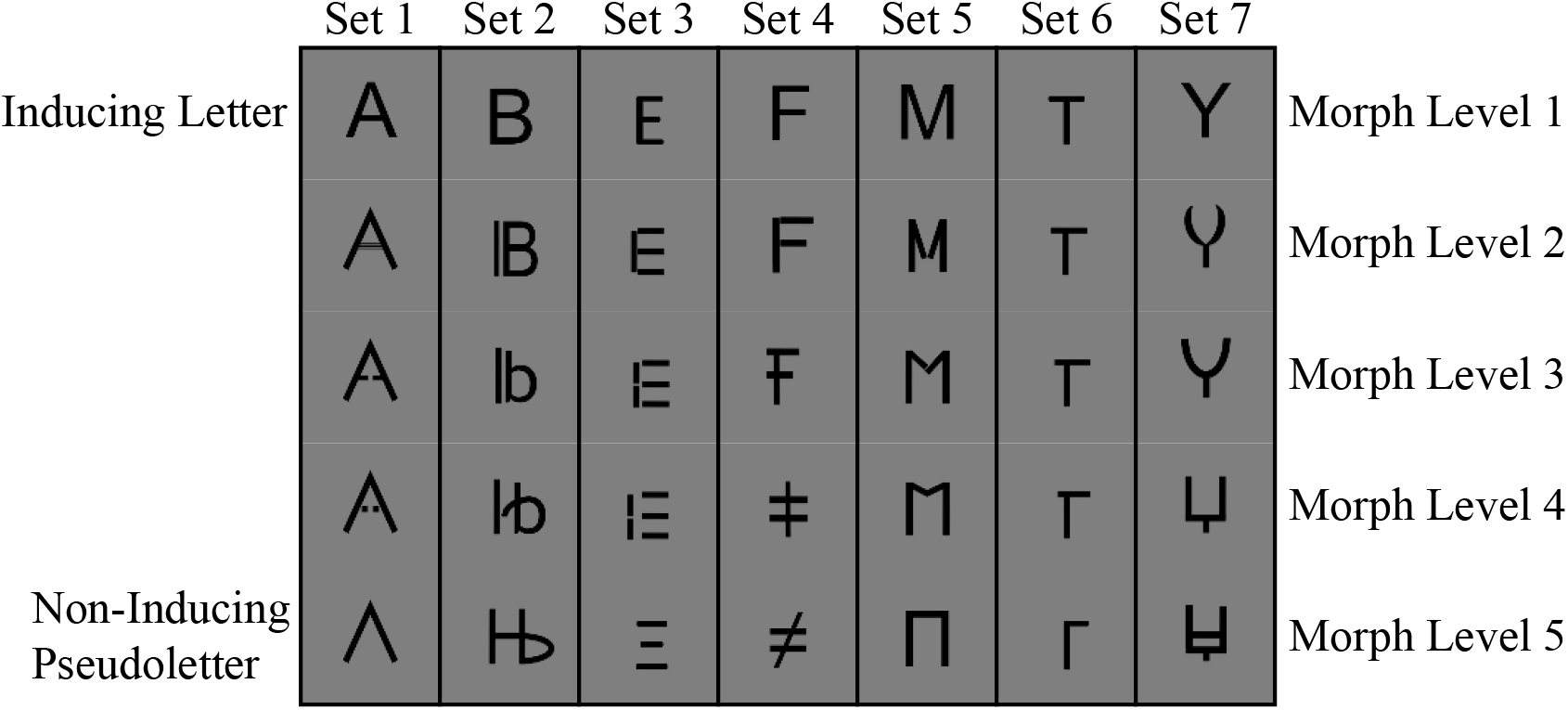
Morph Sets. Seven static sequences of morph graphemes were created for each grapheme-colour synaesthete, such that each complete “morph set” consisted of a colour-inducing letter, a non-inducing pseudo-letter, and three “intermediate” morph graphemes representing step-wise transformations between the inducing letter and the non-inducing pseudo-letter. The intermediate morph graphemes were created such that they physically resembled a “blend” of their preceding and subsequent graphemes.

### 2.4 Psychophysics of the Synaesthesia-Inducing Stimuli

Following creation of the stimulus sets, all synaesthetes (but not controls) were asked to complete a computer task aimed at: 1) acquiring psychophysical measures of the synaesthesia-inducing and non-inducing nature of the presented stimuli across the five morph levels, and 2) defining the optimal duration of stimulus presentation for the subsequent MEG task (i.e. to minimize stimulus duration without compromising synaesthesia induction for synaesthesia-inducing graphemes).

Task (see also Figure 3): Participants were instructed to focus their attention to the centre of the screen. Each trial began with the presentation of an instruction screen (medium grey background) prompting the participant to press the “spacebar” key when ready for the next trial. Upon the key-press, the stimulus appeared in black against the medium grey background. The stimulus was always one of the thirty-five pre-selected graphemes (i.e., from that particular synaesthete’s stimulus set). The stimulus remained on the screen for a pre-defined stimulus duration time (either 50 ms, 200 ms, or 1000 ms, randomized across trials). Fifteen repetitions were presented per stimulus and presentation time, which resulted in a total of 525 trials.

**Figure 3.**
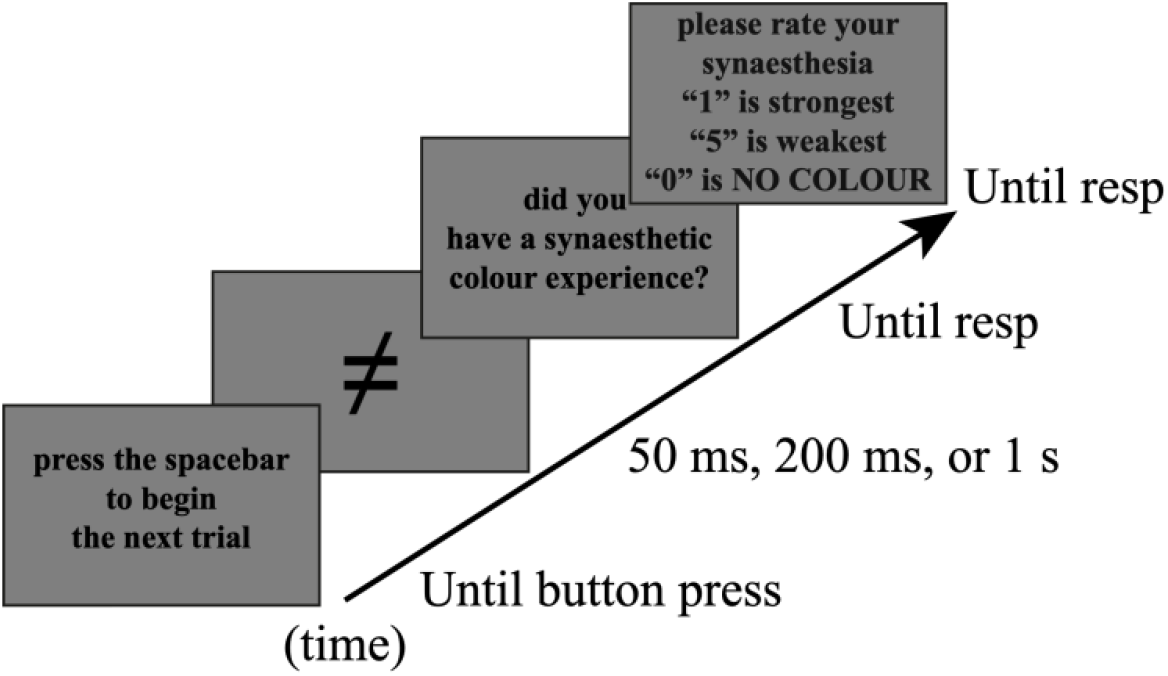
Task used in Psychophysical Testing of MEG Stimuli.

The task was two-fold. Synaesthetic participants were prompted first to indicate by button press whether the presented stimulus had induced a synaesthetic colour experience or not, by using two pre-defined (yes-no) keys; and second to rate, on a scale from 0 to 5, how strong their synaesthetic experience was via six other pre-defined keys (marked with labels 0-5). Synaesthetes were instructed to always respond with a ‘0’ if they had *not* experienced a synaesthetic colour, and to rate the strength of their synaesthetic colour experiences from 1 to 5 if they had answered “yes” to the previous question, with “1” being the strongest and “5” being the weakest synaesthetically induced colour experience. Synaesthetes were encouraged to use all five button presses and were reminded in every trial via on-screen instructions of the response-key assignments. Both questions, presented sequentially, remained on the screen until response. Synaesthetic participants were encouraged to take breaks, as the task lasted between 60-90 minutes, depending on individual pace.

### 2.5 MEG Task

During MEG recordings, grapheme-colour synaesthetes and matched controls viewed pairs of achromatic graphemes presented sequentially, while performing a grapheme comparison task of the two (Figure 4). The aim was to compare cortical processing of synaesthesia-inducing letters to non-inducing pseudoletters. Morphs were also presented in order to maintain participants’ attention to the presented graphemes (i.e., to increase the difficulty of the comparison task), but were not analyzed in terms of MEG responses.

**Figure 4.**
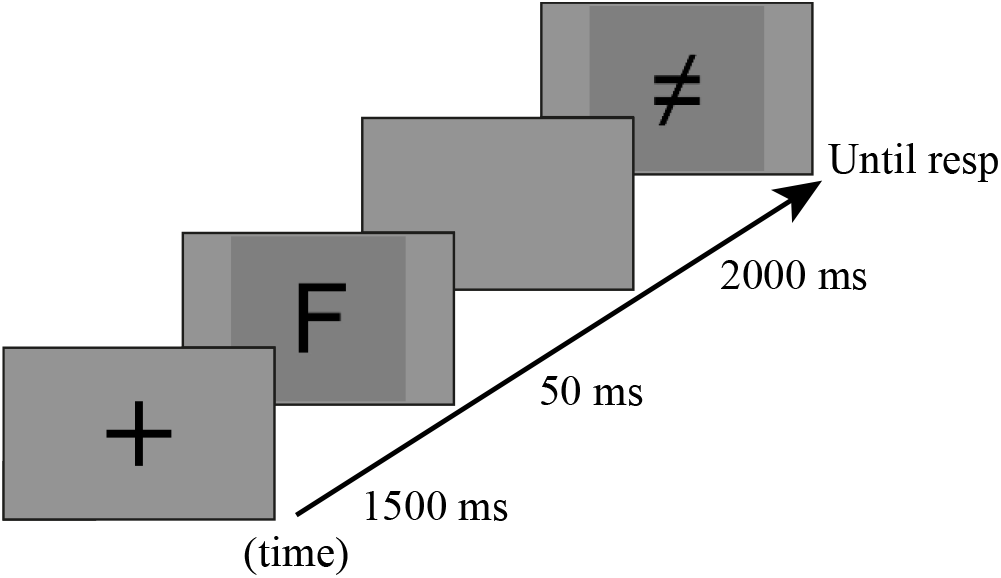
Task administered to grapheme-colour synaesthetes and controls in MEG. Participants were instructed to attend to all stimuli and rate the similarity between presented graphemes (on a scale from 1-5) following the presentation of the second one.

In each trial, synaesthetes and controls viewed two successive achromatic graphemes, drawn from the pool of pre-selected graphemes (i.e., the individual stimulus set). The presented graphemes could thus be colour-inducing letters, morphed graphemes, or non-inducing pseudoletters (i.e., any two graphemes *within* a stimulus set). Participants were instructed to compare the two presented graphemes on a scale from 1 to 5, where ‘1’ was “very similar” and ‘5’ “very different,” and the numbers ‘2,’ ‘3,’ and ‘4’ progressively dissimilar. Participants were encouraged to use all five buttons.

Stimuli were presented through a DLP projector (PT-D7700E-K, Panasonic) placed outside the shielded room onto a screen situated 1.90 m away from the participants via an in-room mirror. All stimuli (achromatic) were presented using Psychtoolbox (Brainard, 1977). Each trial began with the presentation of a black fixation cross against a medium grey background. After a delay of 1.5 s, the first grapheme (stimulus 1) was presented for 50 ms (duration selected based on the psychophysical pre-tests detailed above). After a delay of 2 s (with only the background screen remaining on the display), a second (different) grapheme (stimulus 2) was presented and remained until response. Upon response (using keys numbered 1-5), the background screen was again presented for 1 s before the next trial began (i.e., signalled by the presentation of a fixation cross). The fixation cross and graphemes were presented in the centre of the screen, and subtended a visual angle of 6 and 4 degrees, respectively.

Each grapheme was presented a total of twelve times as stimulus 1, and was paired with each of the other four related graphemes in its stimulus set three times (stimulus 2, separated by a blank screen, as described above). This led to a total of 84 trials (7 stimulus sets x 12 repetitions) per each of the five morph levels (synaesthesia-inducing letter, morph 2, morph 3, morph 4, non-inducing pseudoletter), and thus 420 trials in total (84 trials x 5 morph levels per stimulus set). Stimulus presentation was divided into six blocks, lasting 6-8 minutes each.

Participants were given instructions to maintain a steady gaze at the centre of the screen, and to blink immediately upon response. They were given unlimited time to rest between runs. On average, the total duration of the task was ∼1 h.

### 2.6 MEG Recording

Brain activity was recorded with a 248-magnetometers whole-head MEG system (MAGNES^®^ 3600 WH, 4-D Neuroimaging) confined in a magnetically shielded room. MEG signal was acquired at a 1017 Hz sampling rate.

Before starting the recording session, 5 coils were positioned on the participant’s head, which was localized at the beginning and end of each run. These coils, together with 3 fiducial points and the subject’s head shape, were digitized using a Polhemus system. During the recording session, participants were seated in a reclining chair and supported their head against the back and top of the magnetometer. Participants were asked to remain as still as possible and were continuously monitored by video camera. They were also instructed to minimize blinking during the presentation of visual stimuli, and instead to synchronize their blinks with the blank grey screen that immediately followed their response.

### 2.7 MEG Analysis

The analysis of the MEG signal was performed using the FieldTrip software package (Oostenveld, Fries, Maris, & Schoffelen, 2011) (see http://fieldtrip.fcdonders.nl/) and in-house Matlab code. It was performed in four main steps: 1) preprocessing aimed at removing artifactual activity; 2) an Independent Component Analysis (ICA) aimed at extracting the dominant patterns of brain activity; followed by 3) a Cluster-Level Analysis on the resulting event related fields (derived from single ICs) evoked by the synaesthesia-inducing (vs. non-inducing) visual stimuli; and, finally, 4) source-level analysis aimed at projecting single ICs into source space, and thus identifying the neural generators underlying the differences between conditions (inducing vs. non-inducing).

#### 2.7.1 Preprocessing

Signals were first epoched in trials of 3 s length (1 s pre-stimulus), time-locked to the onset of the first stimulus in the pair (stimulus 1). We then removed the DC offset and linear trends in the signal to centre it around zero. To standardize the whole-signal preprocessing and facilitate subsequent source analysis, a common set of MEG sensors (n=8) manifesting low correlation with immediate neighbours (signifying increased levels of hardware noise) were removed from the MEG data set. These MEG sensors were manually selected by computing the correlation between individual channels and their first and second order neighbours over the entire signal length (with bad trials removed, i.e., trials manifesting a variance three z-scores above the average variance, per channel). Then, trials contaminated with SQUID jumps were discarded from further analysis, and the remaining MEG signal was de-noised relative to the MEG reference sensors, as implemented in the “ft_denoise_pca” function in FieldTrip. Finally, trials with large signal variance were removed from the MEG data set prior to implementing ICA to isolate and reject both eye blinks and cardiac components from the MEG signal (“fastica” algorithm implemented in FieldTrip, after a dimensionality reduction to 20 components).

#### 2.7.2 Independent Component Analysis (ICA) for Analysis of Evoked Signals

In the case of comparing two experimental conditions, as is done here, performing ICA to each of the conditions separately could lead to the undesired situation in which the decomposition of a component was not performed in exactly the same numerical way for both conditions. In such a case, it becomes difficult both to identify and compare components underlying a brain process present in both conditions, but dominant in only one. For these reasons, ICA was performed on the entire data set before isolating the conditions of interest (Inducing vs. Non-Inducing).

Specifically in this study, *ICA has been employed to isolate components present in both conditions of interest*, on a single-subject level. All components are then compared across the conditions of interest (Inducing vs. Non-Inducing Graphemes), in order to identify components dominated by one condition versus the other. Thus, following the preprocessing of the raw data, the “cleaned” data were downsampled to 250 Hz and subjected to an ICA (“runica;” FieldTrip/EEGLAB, http://sccn.ucsd.edu/eeglab/) in a time window between -0.3 s and 1.2 s. This algorithm first performs a PCA-based dimensionality reduction to 40 components, and then performs ICA on these 40 components. For each participant, the resulting data (from the ICA) were filtered between 1-50 Hz, since only event-related averages were of interest; and finally, single trials in each ICA component were averaged separately for both conditions (Inducing vs. Non-Inducing).

### 2.7.3 Nonparametric Cluster-Based Permutation Analysis (ICA Space)

We then applied a nonparametric cluster-based permutation analysis (Maris & Oostenveld, 2007), as implemented in FieldTrip, to the resulting single-subject data (i.e., from each participant’s ICA) in order *to identify clusters of time in which the two conditions of interest (Inducing vs. Non-Inducing) exhibited significant differences* (time window of interest, 70-320 ms). This test controls the family wise error rate (FWER) in the context of multiple comparisons. For each permutation (n=1000), time clusters are defined on the basis of temporal adjacency by regrouping samples whose t-values correspond to (or exceed) a p-value of 0.05. Cluster-level statistics are then calculated by taking the sum of t-values within the cluster. Here, only temporal clusters with corrected p-values ≤ 0.025 are reported (note that the 97.5^th^ quantile corresponds to the threshold for a two-sided parametric t-test at critical alpha-level 0.05, as was performed here). For each parcitipant, only ICs surviving the cluster-based permutation analysis (p<0.05) were kept and further examined (timing of significant differences, inverse solution for individual ICs).

### 2.7.4 Timing of Significant Differences between Inducing and Non-Inducing Conditions within Independent Components (ICs)

In order to further refine the selection of significant ICs across participants, the time window of maximal temporal overlap across participants’ significant clusters was identified. Importantly, this allowed identification of the time period in which processing of the Inducing grapheme differed from that of the Non-Inducing grapheme across all participants. To this end, all ICs exhibiting significant differences between conditions were grouped together independently for each group (Synaesthetes vs. Controls), and the distribution of significant time points (i.e., time points corresponding to significant differences between conditions) was plotted in time-bins of 20 ms. The time window corresponding to maximum temporal overlap (across participants’ significant clusters) was thus identified, and only corresponding ICs were further analysed (i.e., those containing significant clusters at least partially falling within the identified time window).

#### 2.7.5 Topography of Significant Independent Components (ICs)

The grand average of these ICs was then calculated individually for each group by projecting ICs back to sensor space on a single-subject level. Planar gradient magnitudes were then computed considering first- and second-order neighbouring sensors (maximum distance of 7.4 cm) using the “sincos” approach implemented in FieldTrip, before the resulting data were averaged across participants of each group. The aim here was to identify and compare the average brain activity and topography in this (highly significant) time period across individual participants of each group, and also between groups.

#### 2.7.6 Source Level Analysis

The ICs (for each participant) showing significant differences between conditions were localized in source space using a weighted-Minimum Norm Least Squares Estimation (wMNLS). The brain source space was created by constructing a semi-realistic single shell head model (Nolte, 2003) from each participant’s own MRI image.

#### 2.7.7 MEG-Magnetic Resonance Image Co-Registration

T1-weighted structural magnetic resonance images (MRIs) of each participant were co-registered to the MEG coordinate system by a semi-automatic procedure that provided the best fit between the participant’s scalp surface, extracted from his/her anatomical MRI, and the digitized head shape from the MEG. To obtain a first approximate alignment between MEG and MRI coordinates, we manually located the three digitized fiducial points (nasion, left and right pre-auricular points) in each individual’s MRI.

#### 2.7.8 Head and Forward Models

The brain was segmented using the segmentation routine implemented in FieldTrip/SPM8 (http://www.fil.ion.ucl.ac.uk/spm). Cortical surfaces were first extracted with the FreeSurfer image analysis suite, which is documented and freely available for download online (http://surfer.nmr.mgh.harvard.edu/; Reuter et al., 2012). Then, the source space spanning the cortical sheet was created using the MNE-suite software (Gramfort et al., 2013; Dale et al., 1999), which by using the topology of a recursively sub-divided icosahedron on the cortical surface, inflated to a sphere, selects the subset of vertices that define the source space. In this work, the original cortical sheet point set from the Freesurfer segmentation was downsampled to a total of 8,196 vertices for each individual. We then constructed a semi-realistic single shell head model (Nolte, 2003) based on each individual’s brain. Finally, we computed the lead fields corresponding to the 2 tangential orientations for each voxel.

#### 2.7.9 Inverse Solution (Source Space)

The aim of the inverse solution, as used here, is to project single ICA activity into source space using a methodology similar to that previously applied to resting state MEG data (de Pasquale et al., 2010; Mantini et al., 2011), in which a weighted-Minimum Norm Least Squares Estimation (wMNLS) is employed (Lin et al., 2004), but with a different regularization parameter for each IC. In particular, each map of the IC spatial weights is projected from sensor to source space through wMNLS and the regularisation parameter is computed based on the distribution of the IC weights (for details, please see Appendix). The inverse solution was computed in MATLAB using the Fieldtrip toolbox (Oostenveld et al., 2011).

## 3. Results

### 3.1 Psychophysics of the Synaesthesia-Inducing Stimuli

As expected, the proportion of trials in which synaesthetes indicated a synaethetic experience decreased across the 5 five morph levels, with optimal duration of stimulus presentation being 50 ms (i.e. minimum stimulus duration without compromising synaesthesia induction for synaesthesia-inducing graphemes). (grapheme-colour synaesthetes collapsed; stimulus duration time = 50 ms; letters, morph 2, morph 3, morph 4, pseudoletter: 91.43 % ± 5.47, 90.95 % ± 5.49, 84.76 % ± 7.18 %, 66.19 % ± 11.10, 40.95 % ± 12.96), as did the mean strength of subjective synaesthesia-experience (max=5, min=0) (3.43 ± 0.54, 3.03 ± 0.48, 2.54 ± 0.42, 1.79 ± 0.433, 0.82 ± 0.32). Comparing the strength of synaesthetic sensations between letters and pseudoletters for synaesthetic participants (n=6) (group-level paired samples t-test) revealed a significant difference (t(5)=4.62, p=0.005), confirming that letters did indeed induce a synaesthetic experience, while pseudoletters either did not (synaesthetic strength=0) or only did so very weakly, as intended per design.

### 3.2 Non-parametric Cluster-Level Permutation Analysis on ICs

The single-subject level ICA on the cleaned, raw signal yielded 40 independent components (ICs) per participant, of which on average 1.6 ICs in Synaesthetes proved to show significant differences between synaesthesia-inducing and non-inducing stimuli (vs. on average 1 IC in Controls), according to a non-parametric cluster-level permutation analysis. Therefore, Synaesthetes generally exhibited more significant ICs than Controls, with all Synaesthetes (but not all Controls) exhibiting at least one significant IC (compare 10 total significant ICs observed in all 6 Synaesthetes vs. 6 total significant ICs observed in only 4 out of 6 Controls) (Figures 5 and 6).

**Figure 5.**
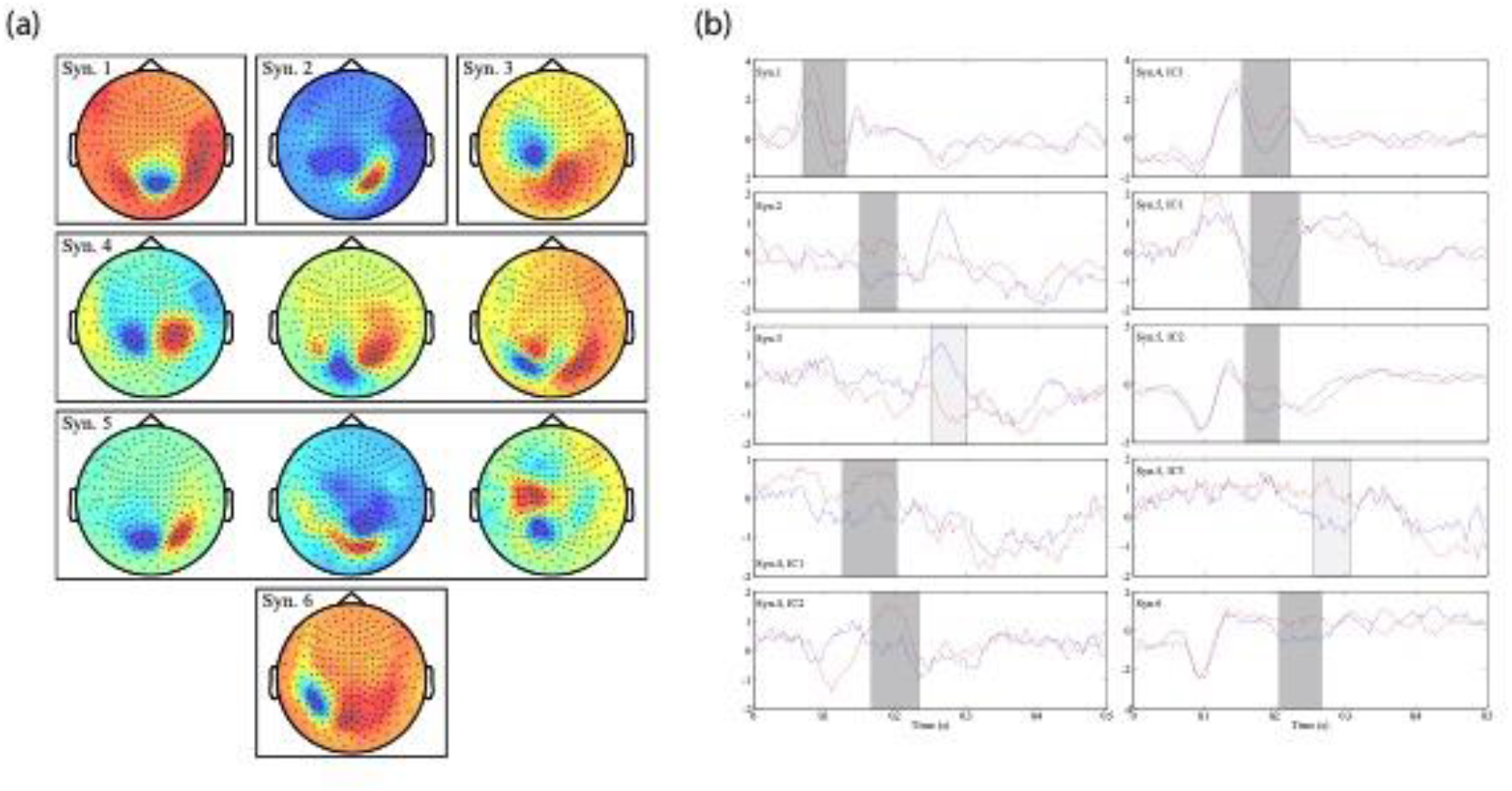
Independent Components of Synaesthetes: Topographies and Time-Series.(a) Represented are the topographies of those ICs exhibiting clusters with significant differences between conditions (Inducing, Non-Inducing) in Synaesthetic participants. (b) Represented are the time series of the same ICs. The shaded areas represent clusters of time in which significant differences between conditions (Inducing, Non-Inducing) occurred. The lighter shading corresponds to ICs with time periods outside the window of maximal temporal overlap (see Figure 7). Note that values on the y-axis correspond to an order of 10e-12 (arbitrary units).

**Figure 6.**
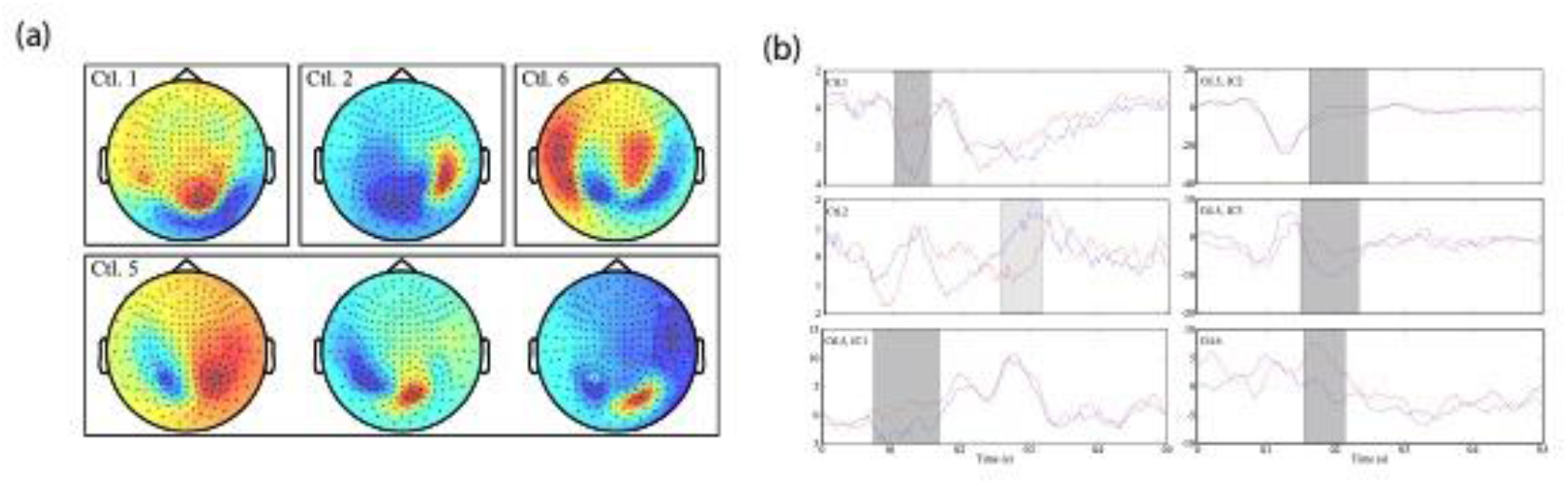
Independent Components of Controls: Topographies and Time-Series. As Figure 5, but for Control participants.

### 3.3 Timing of Significant Differences between Inducing and Non-Inducing Conditions within Independent Components (ICs)

In order to identify the time window of maximal temporal overlap across participants’ significant clusters (i.e., the time period in which significant differences between conditions generally manifested across participants), all ICs containing significant clusters (i.e., significant differences between conditions) were grouped together (independently for each group, Synaesthetes vs. Controls). First, the entire analysed time window (70-320 ms) was divided into bins of 20 ms. Then, the time points of significant clusters were grouped into the specified time-bins (i.e., 70-90 ms, 90-110 ms, 110-130 ms, etc.). The total number of significant clusters falling within each time-bin was counted, normalised according to the total number of participants in each group (n=6 for each), and plotted (Figure 7).

**Figure 7.**
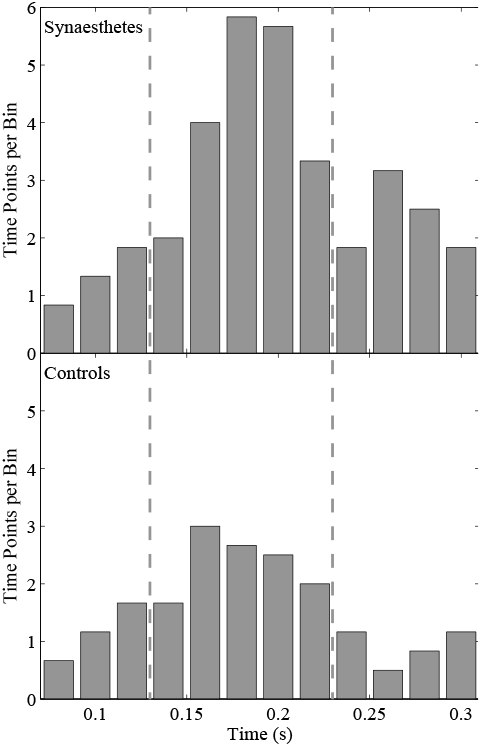
Maximal Temporal Overlap of ICs. Bar plots for each group (Synaesthetes, Controls) showing the frequency of significant clusters (within ICs) for the entire analysed time window (70-330 ms), in bins of 20 ms and normalised to number of participants in each group (n=6).

Figure 7 thus illustrates the number of clusters (within ICs) that show significant differences between Inducing and Non-Inducing graphemes across the analysed time window (70-320 ms), in bins of 20 ms, for each group. We report three main findings from this analysis. First, significant differences in processing of Inducing and Non-Inducing graphemes occur in synaesthetes predominantly in a late time window, peaking at around 190 ms (ranging between 130-230ms). Second, the histogram demonstrates that synaesthetes show more significant differences than controls. Third, controls showed a similar timing for differences between both experimental conditions but with fewer ICs.

### 3.4 Topography of Significant Independent Components (ICs)

Since the maximum temporal overlap of significant clusters across participants in both groups centred at approximately 190 ms and the majority of clusters fell within 130-230ms, the grand average of all corresponding ICs (i.e., those containing significant clusters at least partially falling within the time window, 130-230 ms) was calculated individually for each group, in order to identify and compare the average brain activity and topography in this (highly significant) time period across individual participants of each group. Figure 8 illustrates the topographies and signal differences of Synaesthetes and Controls between conditions (Inducing vs. Non-Inducing) within the pre-selected time window (130-230 ms). While the topographies in panel (a) show the contrast between conditions (Inducing minus Non-Inducing) for each group independently (Synaesthetes and Controls), the topography in panel (b) shows the difference between these (Synaesthetes minus Controls), revealing increased activity in Synaesthetes (vs. Controls) in occipito-parietal areas.

**Figure 8.**
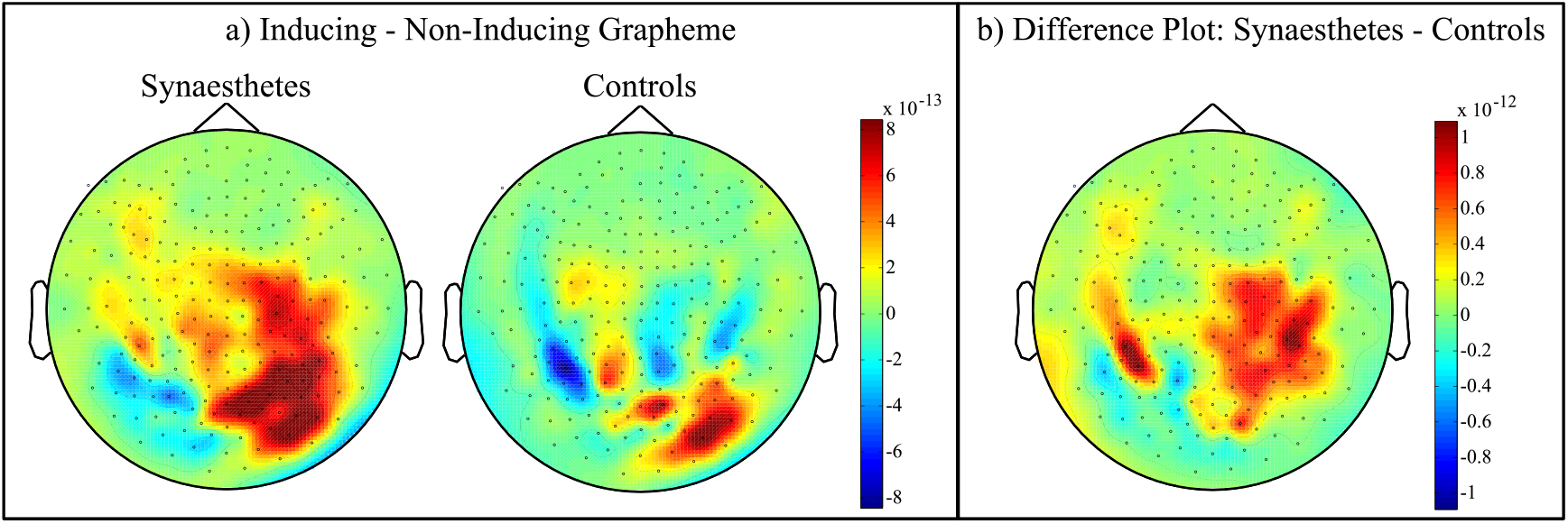
Contrast between Inducing and Non-Inducing Conditions. The topographies and signal differences between conditions (Inducing vs. Non-Inducing) within the pre-selected time window (130-230 ms) are shown. (a) Contrast between conditions (Inducing minus Non-Inducing) for each group independently (Synaesthetes, Controls). (b) Contrast between groups (Synaesthetes minus Controls). Note that this difference plot is a difference (Synaesthetes minus Controls) of two differences (Inducing minus Non-Inducing, for each group).

### 3.5 Source Level

The wMNLS source reconstruction, performed on a single-subject level, yielded consistent sources across the five Synaesthetes (of 6) who showed ICs in the 130-230 ms window (n=5, see Figure 9). As indexed by the Talairach Tournoux atlas (Talairach & Tournoux, 1988), these were localized to visual extrastriate cortex overlapping with Brodmann area 19 in the occipital lobe (in five Synaesthetes) and Brodmann area 7 in the superior parietal lobe (in three Synaesthetes). In contrast, source reconstructions across the three Controls (of 6) who showed ICs in the 130-230 ms window were less consistent (see Figure 10), localizing instead to Brodmann area 18 in the occipital lobe in one participant, to Brodmann area 40 in the inferior parietal lobe in a second participant, and to both areas in a third.

**Figure 9.**
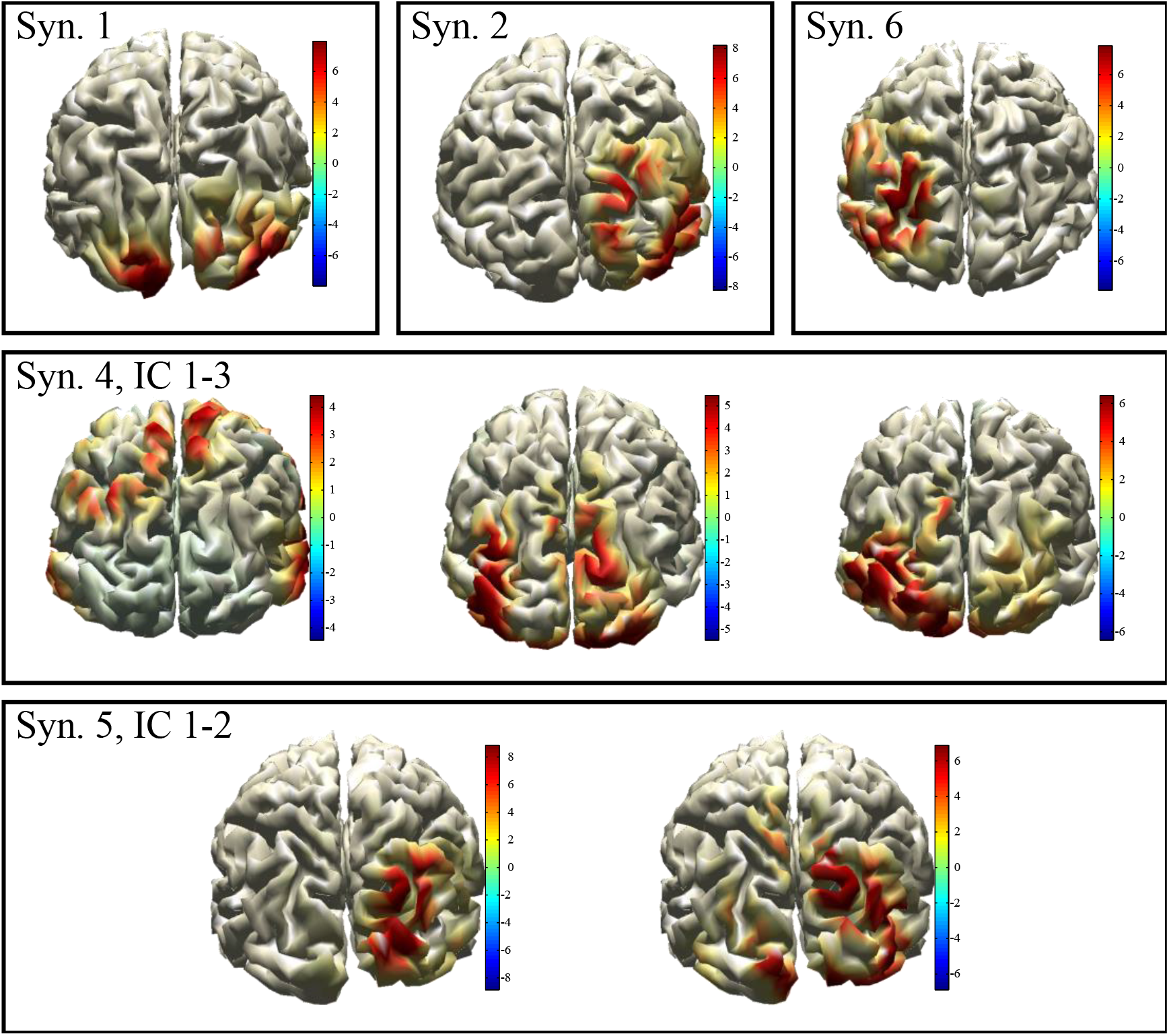
wMNLS source reconstructions in individual Synaesthetes. Five out of six Synaesthetes exhibited ICs with significant differences between conditions. The inverse solutions of these individual ICs are shown. All Synaesthetes but one (Syn.1) exhibited sources both in the occipital and parietal lobes.

**Figure 10.**
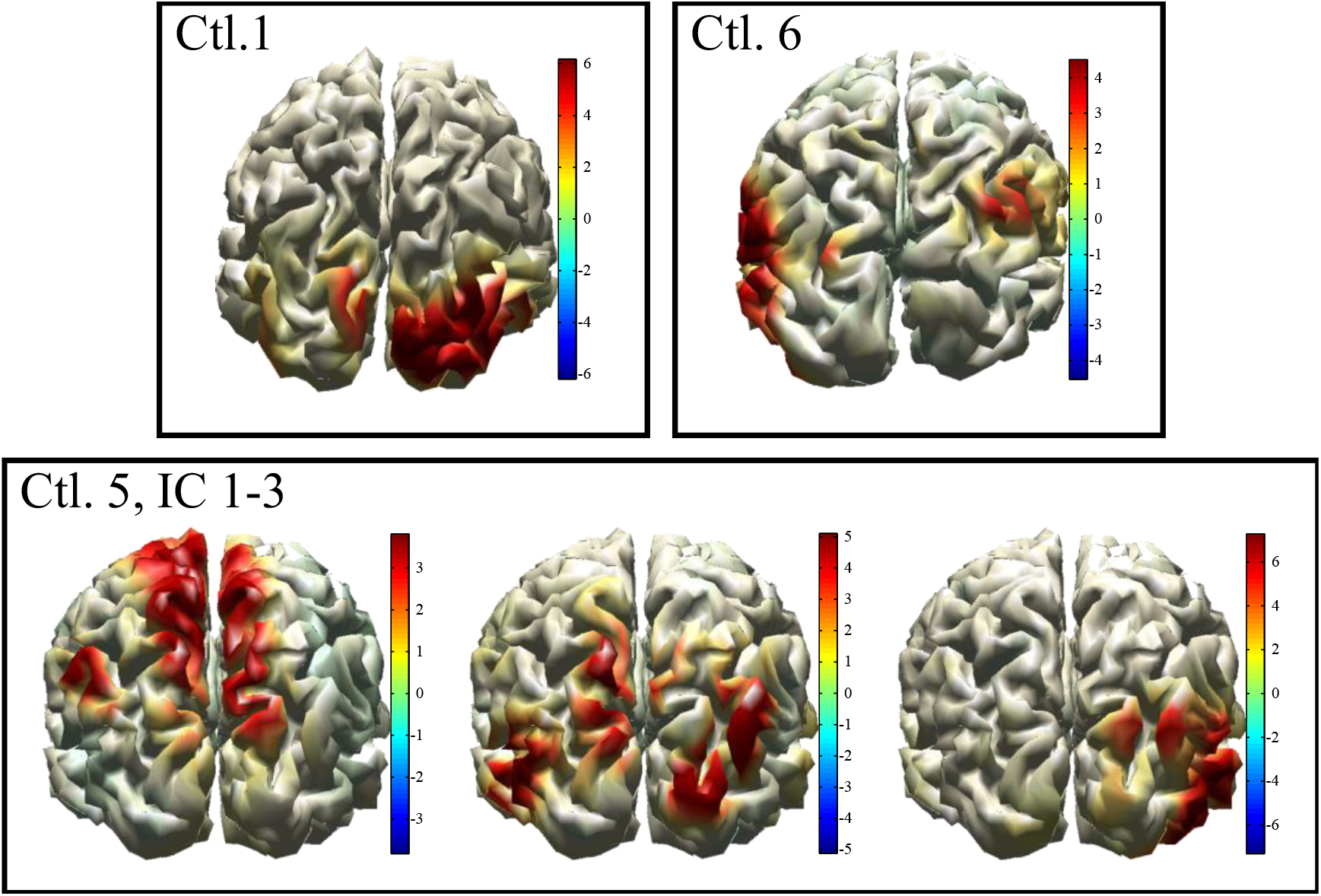
wMNLS source reconstructions in individual Controls. As in Figure 9, but for Controls. In contrast to Synaesthetes, sources across the three Control participants were inconsistent, localizing to differing occipital, temporal, and parietal areas.

Note that the scaling of Figures 9 and 10 reflect Z-scores of the projected IC weights. What are projected through the inverse solution into source space are the weights of single ICs. The spatial filters are derived through wMNLS, which uses the forward solution as a model for propagation (from the brain to the sensors). This inverse solution translates the magnetic field measurements outside the brain into activation inside the brain. In our case, we multiply the unit-less set of numbers (i.e., weights of ICs) with this inverse solution; consequently, these numbers (i.e., the weights of the ICs) are scaled by the spatial filter of the inverse solution. As the IC weights are unit-less, this scaling does not produce a physical quantity inside the brain (such as power), but rather a set of linearly combined, scaled weights. In order to make these results more interpretable, we add a further scaling by using the Z-score of the projected numbers. This further scaling provides a better interpretation of the distribution of the projected weights on the cortical sheet.

## 4. Discussion

We here carry out an MEG study on associator grapheme-colour synaesthetes in whom we examine the neural activity dissociating the perception of synaesthesia-inducing letters from non-inducing pseudoletters. Overall, we find that the neural processes differentiating between inducing and non-inducing graphemes occurs relatively late in the processing hierarchy, peaking approximately at 180-200 ms, well after grapheme identification is likely complete (Rey, Dufau, Massol, & Grainger, 2009) and consistent with a recent MEG study also examining grapheme-colour synaesthetes (Teichmann et al., 2021). Due to the late timing of these effects, they do not likely occur during the initial, feed-forward sweep of activity in the visual processing stream (see Lamme & Roelfsema, 2000), as predicted by models of synaesthesia invoking rapid timing predictions (i.e., Cross-Activation/CCT models). In addition to the timing of induced synaesthetic activity, we performed source reconstruction of the corresponding signals and report consistent involvement of occipito-parietal areas, localising consistently to extrastriate visual cortex in the occipital lobe and coinciding with activity in the superior parietal lobules. These anatomical correlates could implicate multisensory convergence zones in the dissociation between synaesthesia-inducing and non-inducing graphemes. Overall, due to both the relatively late timing as well as anatomy of these neural events, our results are more consistent with models of synaesthesia supporting a relatively late synaesthetic correlate, such as Disinhibited Feedback or any of its variants (Grossenbacher & Lovelace, 2001).

There are few electrophysiological studies examining the underlying mechanisms of induced, synaesthetic percepts in grapheme-colour synaesthesia. Although several of these studies report early processing differences between synaesthetes and controls beginning as early as 100 ms after viewing graphemes (Brang et al., 2008; Brang et al., 2011) or hearing words (Beeli et al., 2005), they primarily addressed *semantic* modulation of synaesthesia-inducing graphemes (i.e., a phrase followed by a synaesthesia-inducing stimulus, whose induced colour is semantically congruent or incongruent to the expected meaning). In addition, several of these studies could not accurately localise the underlying electrophysiological generators due to volume conduction limitations characteristic of EEG data. Similarly, the only MEG study to-date examining the anatomical underpinnings as well as the timing of grapheme-colour synaesthesia (Brang et al. (2010)) showed early (i.e., beginning around 110 ms) activity in area hV4 in response to synaesthesia-inducing graphemes in synaesthetes (versus controls), only ∼5 ms after the onset of activity in adjacent grapheme processing areas. The authors concluded that this short latency (∼5 ms) between activity in grapheme processing areas and hV4 could only be supported by direct, anatomical connections between the two areas, as predicted by Cross-Activation/CCT models. However, the four grapheme-colour synaesthetes included in the study by Brang et al (2010) were strong *projector* synaesthetes, and thus it is unclear whether their results may generalise to the majority of synaesthetes, who are associator sub-types. Associators differ from projectors in that their synaesthetic qualia are more conceptual than perceptual and are generally experienced in their mind’s eye rather than in external space. It is thus possible that different models account for different synaesthetic sub-types, given their differences in phenomenology, measured behaviour, and differential white matter connectivity (Rouw et al., 2010; Rouw et al., 2007).

The difference topography between Synaesthetes and Controls (derived from the group averages of all ICs exhibiting significant differences between conditions) implicates occipito-parietal areas (see Figure 8) peaking at ∼190 ms. Furthermore, localisation of significant ICs via a Minimum Norm inverse solution (single-subject approach) consistently yielded areas in extrastriate occipital cortex and both inferior and superior parietal lobes (Figure 9). The superior parietal lobes have been implicated in the (spatial) co-localization of visual features into coherent percepts (i.e., feature conjunction tasks) (Baumgartner, 2013; Robertson, 2003; Donner et al., 2002; Shafritz et al., 2002; Corbetta et al., 1995) and, importantly, have been causally implicated in the binding of colours to letters in grapheme-colour synaesthesia (Esterman, Verstynen, Ivry, & Robertson, 2006; Muggleton, Tsakanikos, Walsh, & Ward, 2007; Rothen, Nyffeler, von Wartburg, Muri, & Meier, 2010). In fact, there is increasing evidence showing the importance of parietal cortex in associator grapheme-colour synaesthesia, particularly the superior parietal and intraparietal sulcus (IPS) regions (van Leeuwen et al., 2010; Weiss et al., 2005; Zeki & Marini, 1998), including anatomical studies showing increased coherence (FA) in the white matter of IPS (Rouw & Scholte, 2007), and functional connectivity studies (Jancke & Langer, 2011; Specht & Laeng, 2011) demonstrating important hubs in parietal areas (in addition to corresponding early sensory areas, such as fusiform gyrus). Thus, parietal cortex seems to play a crucial (essential) role in the induced synaesthetic percept of associator synaesthetes, either in the hyperbinding of visual features elicited in earlier visual areas, or in feedback to earlier visual areas.

Although lateralisation of occipital and parietal areas varied between Synaesthetes, previous neuroimaging studies have reported similar findings (see (Gray et al., 2006; Zeki & Marini, 1998)). In fact, synaesthetic colour has been found to activate a broad range of areas in ventral occipitotemporal cortex (see Rouw et al., 2011, for a review). Synaesthesia is highly idiosyncratic and individual differences between synaesthetes are common in the literature, both in behaviour and neuroimaging (Rouw & Scholte, 2010; Sperling et al., 2006; Hubbard et al., 2005; Dixon et al., 2004), including lateralisation.

Contrary to Synaesthetes, the three Controls who also displayed significant differences between conditions showed no consistency in their inverse solutions (Figure 10), possibly reflecting task-specific strategies, slight differences in activity reflecting letters versus pseudoletters, or possibly further supporting a model of synaesthesia invoking enhanced activity in universally-present neural pathways (i.e., akin to Stochastic Resonance). We did not expect to observe early visual differences between conditions (Inducing vs. Non-Inducing) based solely on grapheme recognition, given (1) the physical similarity between letters and pseudoletters in terms of low-level visual complexity, (2) current theories of grapheme recognition as a process of hierarchical feature analysis, and (3) the lack of letter-centred or language-centred task demands (note that 80% of presented graphemes were pseudoletters, or morphed graphemes) (Dehaene et al., 2005; Hubbard et al., 2005; Mitra & Coch, 2009) (but see Rey et al., 2009). Nonetheless, we cannot rule out the possibility that observed differences between conditions reflect visual processing differences unique to grapheme-colour synaesthesia, but not specifically related to induced, conscious colour concurrents. Since our paradigm did not directly address the “consciousness” or phenomenological experience of the synaesthetic concurrent itself, these are here not dissociable from general processing differences between inducing and non-inducing stimuli.

We here present evidence for the timing of induced activity in associator grapheme-colour synaesthetes, more in line with models like the Disinhibited Feedback model, implicating functional (as opposed to structural) differences between synaesthetes and non-synaesthetes. This is consistent with models proposing common mechanisms of cross-modal integration across synaesthetes and non-synaesthetes alike. In sum, our study is the first MEG study to date revealing stimulus evoked activity in associator grapheme-colour synaesthetes in a network of areas including the occipital and parietal cortices, peaking relatively late (∼190 ms) in the visual processing stream of events and thus advancing our ability to disentangle between current models of synaesthesia.

## Appendix

### Minimum Norm Inverse Solution for Single Independent Components

The weighted Minimum Norm Least Squares (wMNLS) solution is computed according to (see Lin et al., 2004):

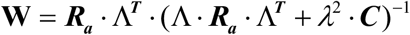

**where**

Λ : Leadfield matrix

***R***_***a***_ : a priori assumed brain source covariance

***C*** : noise covariance in MEG sensor array

*λ* : Minimum Norm regularisation parameter

In our work, we followed a novel approach for the computation of the regularization parameter for each IC, according to the following equation:

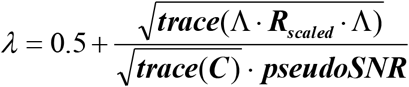

**where *R***_***scaled***_ is the a priori assumed brain source covariance matrix scaled as :

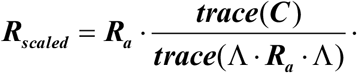

so that

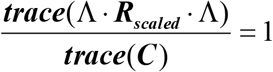

Consequently, the regularization parameter formula is reduced to:

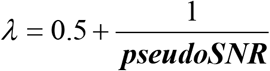

and the inverse solution becomes:

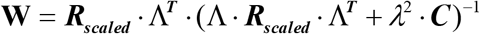

where ***pseudoSNR*** is a scalar parameter used to represent a pseudo Signal-to-Noise ratio for a single IC.

As the noise power within a single IC is unknown, here we chose to derive an empirical measure of how well an ICA represents a few strong focal brain sources or widely distributed noise. For an ICA representing a strong focal brain dipole, the squared ICA unmixing weights have a skewed distribution, with high values at the sensors close to the underlying sources, and all the rest of the sensors (further from the underlying sources) having much lower values. In the case of an IC capturing widely distributed noise, the squared ICA unmixing weights have more comparable values. Consequently, the upper, i.e. 70 %, and lower, i.e. 30 %, distribution percentiles are expected to be more distant in the case of a brain activity IC than in the case of a noise IC.

This parameterization has been used in order to estimate a pseudo Signal-to-Noise Ratio for a single IC. If the squared unmixing ICA coefficients for a single ICA are represented by ***Uic***, then the ***pseudoSNR*** is computed as:

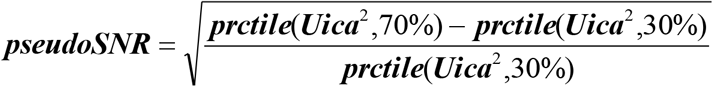

This parameter has a lower bound of 0. The higher the distance between the percentiles, the higher the value of this parameter. The closer the upper and lower percentiles get, the closer this parameter is to this lower bound.

From the formula for the regularisation parameter, the latter term 1/ ***pseudoSNR*** varies in an inverse fashion, from 0 to high positive values. This means that for ICs representing strong dipolar sources, little regularization is used, as the unmixing matrix contains a clear dipole representation. For ICs representing noise, a higher regularization is used as the unmixing matrix represents a more complex and distributed pattern.

Having very small regularisation values close to 0 for very strong dipoles can lead to instability in the derivation of the inverse solution. In order to avoid such instabilities, a scalar value of 0.5 has been added to 1/ ***pseudoSNR*** in the derivation of the regularization parameter. This value represents the 1/ ***pseudoSNR*** ratio when the difference ratio between the upper and lower percentiles under the square root in ***pseudoSNR*** is equal to 4.

With this final formulation, the regularisation parameter varies between 0.5 (for ICs representing strong brain sources) and infinity (for ICs representing noise). Infinity here just represents very high values. This is because in ICA unmixing matrices, the 30 % and 70 % percentiles cannot have the exact same values, as this would require that all the in between weights in the distribution should be identical.

The above described regularisation parameter has a lower bound, which hedges against instabilities of the inverse solution, and no upper bound, which allows for high regularisation when ICA components representing noise are localised.

The above described inverse solution procedure was applied to each of the ICs, for which a significant statistical difference was found in the comparison between the compared conditions, both for synaesthetes and controls. No a priori brain sources covariance was assumed, so **R**_***a***_was the identity matrix with dimensions Nsources x Nsources. As the level of noise in the single ICs was also unknown, the noise covariance matrix **C** was the identity matrix as well, with dimensions Nsensors x Nsensors. The source localization was performed and plotted on the 3-dimentional template grid with 6mm resolution, warped to each subject’s brain volume.

### Figures & Figure Legends

